# Plasticity in parental effects confers rapid larval thermal tolerance in *Nematostella vectensis*

**DOI:** 10.1101/2020.06.15.153148

**Authors:** Hanny E. Rivera, Cheng-Yi Chen, Matthew C. Gibson, Ann M. Tarrant

## Abstract

Parental effects can prepare offspring for different environments and facilitate survival across generations. We exposed parental populations of the estuarine anemone, *Nematostella vectensis*, from Massachusetts to elevated temperatures and quantified larval mortality across a temperature gradient. We find that parental exposure to elevated temperatures results in a consistent increase in larval thermal tolerance (mean ΔLT50: 0.3°C), and larvae from subsequent spawns return to baseline thermal thresholds when parents are returned to normal temperatures. Histological analyses of gametogenesis in females suggests these dynamic shifts in larval thermal tolerance may be facilitated by maternal effects in non-overlapping gametic cohorts. We also compared larvae from North Carolina (a genetically distinct population with higher baseline thermal tolerance) and Massachusetts parents, and found larvae from heat-exposed Massachusetts parents have thermal thresholds comparable to larvae from unexposed North Carolina parents. North Carolina parents also increased larval thermal tolerance under the same high-temperature regime, suggesting plasticity in parental effects is an inherent trait for *N. vectensis*. Overall, we find larval thermal tolerance in *N. vectensis* shows both a strong genetic basis and phenotypic plasticity. Further understanding the mechanisms behind these shifts can elucidate the fate of thermally sensitive ectotherms in a rapidly changing thermal environment.

## Introduction

Parental effects encompass a range of mechanisms aiming to better prepare offspring for the conditions they may experience. These effects are often informed by the parental environment, especially if the environment the offspring will experience will resemble that of the parents. Such anticipatory parental effects allow parents to enhance the phenotypic plasticity of offspring to better match their future environment (1). Parental effects may influence offspring throughout their lifetimes, as for instance, occurs in the water flea *Daphnia*, which will develop a helmeted anti-predation phenotype if parents were exposed to predators (2). Alternatively, effects may be short-lived, mainly influencing early life stages, often through mechanisms such as maternal loading of RNA transcripts; increased energetic reserves (e.g. lipids) in seeds, embryos, and larvae; or modification of the gestational environment, in order to enhance offspring survival or allow for faster acclimatization to environmental conditions (see reviews in 3,4). In an era of rapid climate change, swift phenotypic modifications facilitated by parental effects may become indispensable for species survival (5).

In the context of temperature, the timing and duration of thermal variation relative to reproductive cycles, as well as the persistence of parental effects across future larval cohorts may facilitate the progression from acclimatization to adaptive processes, especially for thermally sensitive organisms (6,7). Transgenerational and parental (often maternal) effects have been shown to promote offspring thermal tolerance in polychaetes (8), fruit flies (9), copepods (10), corals (11), sticklebacks (12), and damselfish (13), among many others. For polychaetes, it appears the impact of maternal effects depends on when during oogenesis the mother experiences a particular environment (8). In *Drosophila*, loading of maternal transcripts for a heat shock protein into eggs, enhanced embryonic thermal tolerance (9). Meanwhile in sticklebacks, maternal effects modulate respiration rates of offspring to promote growth under temperature conditions similar to the mother’s (12). As these results show, parental effects span a wide range of mechanisms and can provide substantial thermal resistance to offspring, facilitating rapid acclimatization to environmental stressors.

The ability of populations to respond to increasing temperatures will play a critical role in determining the distribution and persistence of species as global temperatures rise (14). The adaptive benefit of parental effects will depend on the how reliably parental environments predict the conditions offspring will experience (1,3,15). Organisms that have both short reproductive cycles and multiple reproductive life cycles across their lifetime represent an interesting case study for investigating the role and effectiveness of parental effects in offspring fitness across ecologically relevant timescales.

The estuarine anemone, *Nematostella vectensis* (Stephenson, 1935) inhabits coastal salt marshes along the eastern and western coasts of North America and parts of the United Kingdom, with populations from different locations showing strong genetic divergence (16). *Nematostella vectensis* persists across a wide range of thermal environments across latitudes, and experiences substantial daily and seasonal thermal variation in shallow, estuarine habitats (17). For instance, in Massachusetts, *in situ* seasonal temperatures range from below freezing in winter to above 30°C in summer (17). Populations across latitudes also have different thermal tolerance thresholds during larval, juvenile, and adult stages, with southern individuals exhibiting faster growth and lower mortality at higher temperatures (18). For example, the temperature at which 50% of individuals die (LT50) varies by nearly 2°C between juveniles from Massachusetts and those from South Carolina (18).

*Nematostella vectensis* is also a highly tractable experimental organism for studying development and ecophysiology (19). *N. vectensis* is able to fully regenerate from a small segment, a process that it uses for asexual reproduction and recovery from injury (20), and can be used to generate clonal lineages by bisecting adults (21). Populations of *N. vectensis* can be easily maintained in laboratory conditions and reproductive cycles reliably induced in two- to three-week intervals (20,22). Females eject egg bundles that are fertilized externally by sperm released by males, facilitating controlled crosses (23). While reproductive timing in the wild has not been studied, laboratory animals can be spawned year-round. Given their wide geographic distribution and highly variable habitats, parental effects could serve as a fast and effective way to modulate the thermal limits of larvae during spawning cycles.

Here, we leverage the physiological range of *N. vectensis* and expose parents to an increasing temperature regime during gametogenesis to quantify its influence on parental effects (both maternal and paternal) on larval thermal tolerance, as well as on adult heat tolerance. We test whether parental effects persist through subsequent spawning events and explore possible mechanisms for the induction of larval thermal tolerance. We further compare the impact of parental effects on larval thermal tolerance to differences between locally adapted and genetically distinct *N. vectensis* populations from Massachusetts (MA) and North Carolina (NC). Taken together, our work assesses the relative influence of acclimation, parental effects, and local adaptation in determining thermal thresholds across life history stages.

## Methods

### (a) Animal collection and husbandry

All *Nematostella vectensis* used in experiments were laboratory populations of animals originally collected from the Great Sippewissett Marsh, MA (41.59°N, 70.63°W) or Fort Fisher, NC (33.95°N, 77.93°W). Populations were kept in glass containers under a 12:12 light:dark cycle, and maintained at 18-20°C in filtered natural seawater diluted with deionized water to 15 PSU salinity. Water changes were conducted every two weeks and animals were fed freshly hatched brine shrimp nauplii 4-5 times per week. Animals from Massachusetts (MA) were maintained at the Woods Hole Oceanographic Institution. North Carolina (NC) animals were reared under comparable conditions, until being transferred to MA ∼6 months before experiments (*N* =15 individuals). Animals used for experiments were placed under constant dark conditions at least two weeks prior to the start of experiments, to reduce any confounding effects associated with variability in light levels across treatments. Husbandry of animals used for analysis of gametogenic cycles is described below.

### (b) Histological quantification of gametogenesis in females

This *Nematostella* strain was collected from the Rhode River in Maryland and was kindly provided by Mark Martindale and Craig Magie. Anemones were cultured in 12 PSU artificial seawater (ASW) and acclimated to lab conditions for more than five years (24). Female anemones were maintained at 18°C with ambient light, fed once per month with *Artemia* nauplii until a week before the induction of spawning (spawn day -7). Then, they were fed *Artemia* nauplii daily until one day before spawn induction (spawn day -1). On one day before the induction of spawning, five females representing the “before spawn” time point were anesthetized in 7% (w/v) MgCl_2_ in ASW and then fixed in 4% paraformaldehyde/FSW (v/v) at room temperature for one hour. After an hour, the aboral end was opened and fresh fixative was added for overnight fixation at 4°C. Remaining females were induced to spawn, as described below, except that in the following morning the females were cold-shocked by replacing room temperature culturing ASW with 18°C ASW and placed under normal light conditions. Five females that were observed releasing egg bundles were anesthetized and fixed for the “after spawn” timepoint. After fixation, samples were washed in PBST (phosphate-buffered saline with 0.2% Triton X-100, v/v), five times (10 mins per wash) at room temperature. Samples were then incubated in 1:5,000 diluted SYBR™ Green I (Thermo Fisher Scientific; S7567) and 1:1,000 diluted SiR-Actin (Cytoskeleton, Inc.; CY-SC001) in PBST at 4°C overnight, then washed three times (10 minutes per wash).

Female mesenteries were then dissected out and immersed in modified Scale A2 (4M urea and 80% glycerol in PBS, v/v), and imaged using a Leica Sp5 confocal microscope, with 4 um per z-stack step to quantify the size and number of oocytes before and after spawning. Three mesentery field of views per animal, and 5 animals per treatment (before and after spawn) were imaged and quantified, for a total of 30 views. Oocytes (*N*=496 oocytes: before spawn; 469 - after spawn) were manually circled to quantify area using ImageJ/FIJI imaging software (25,26). The density of oocytes within mesenteries was measured semi-automatically by FIJI macro, in which maximum projected images of F-actin channel (mesenteries) were filtered by Gaussian Blur (Sigma = 8), thresholded with the same value to automatically select oocyte tissue and manually curated to measure the area.

Oocyte data was tested for normality using Shapiro-Wilk tests. As data were non-normally distributed, a ranked two-sample Wilcoxon ranked test was used to quantify differences in oocyte size (area) before and after spawn.

### (c) Establishment of clonal lineages and genotype-controlled parental populations

To control for genetic variability between parents, experiments were conducted with clonal parental populations whenever possible (see individual experimental descriptions below). To create clonal lineages, animals were repeatedly bisected across the body column, and allowed to regenerate completely until lineages reached 20-40 individuals. Multiple individuals from each lineage were combined to incorporate genetic diversity into genotype-controlled parental populations. A detailed enumeration of lineages/individuals in each experimental population is provided in supplementary table 1. NC animals were bisected to obtain a total of 35 female individuals (*N*=3 lineages) and 60 male individuals (*N*=5 lineages).

### (d) Heat stress regime

Using two, Precision™ Dual Chamber 188 water baths (Thermo Fisher Scientific, Waltham, MA, USA), animals were transitioned from 20°C to 33°C, at a rate of ∼3°C/day, held at 33°C for four days (simulating maximum summer noon temperatures in the Sippewissett Marsh; figure 1A), and then returned to 20°C at the same rate, prior to spawning. A HOBO™ Tidbit logger (Onset Computer Corporation, Onset, MA, USA) was used to track treatment temperatures every half hour during experimental incubations. Parental populations exposed to the heat stress regime are hereafter called the short-term heat stress (STHS) parental treatment. Water changes were conducted every other day for both STHS and control animals. Each bowl was fed the same ration of brine shrimp nauplii daily (0.2 grams). Control animals were kept at 20°C in a Low Temperature Incubator (Thermo Fisher Scientific, Waltham, MA, USA), humidified to prevent evaporation. All animals were kept under constant dark conditions.

**Figure 1.**
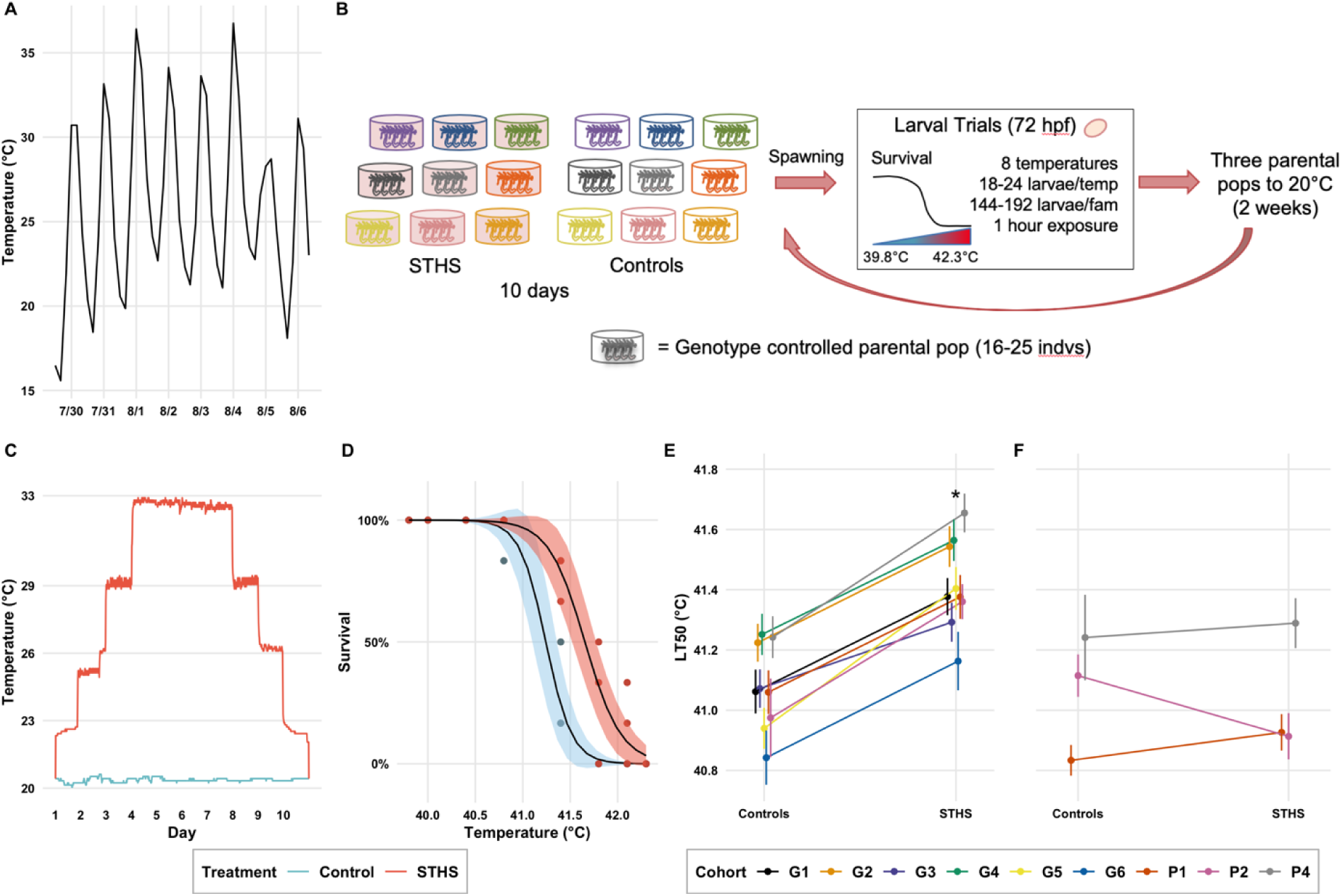
Larval thermal tolerance increases when parents experience heat stress during gametogenesis. (A) *In situ* temperatures in a tidepool containing a *Nematostella vectensis* population within Great Sippewissett Marsh, MA, during one week in summer 2017, logged every four hours. (B) Experimental design for MA parental effects experiment. See table S1 for population breakdown. STHS – Short Term Heat Stress parental treatment (C) Temperature regimes experienced by parents during the 10 days prior to spawning, logged every half hour. (D) Survivorship curves of larvae from control (blue) and STHS (red) parents when exposed to acute temperature stress. Ribbon shows 95% confidence interval for logistic survivorship model. Curve shown for family G6, curves for all families tested in figure S1. (E) Temperature at which 50% of larvae are dead (LT50). Colors correspond to genotype-controlled families. Error bars denote standard error. Mean shift in LT50 is 0.34°C (*p*= 2.25×10^−7^, paired t-test). (F) Larval LT50 for *n*=3 parental pairs that were re-spawned approximately 2 weeks after the first experimental spawn (shown in E). Paired t-test for differences in LT50 between parental treatments were non-significant *p=*0.57, indicating parental effects subside during subsequent spawns.

### (e) Spawning

On the day following the end of the heat stress regime, parental populations were induced to spawn following the protocols detailed in (22). Briefly, *N. vectensis* were individually fed mussel gonad tissue. After 5-6 hours, their water was changed and anemones were placed under a bright full spectrum light for 12 hours overnight at room temperature. The following morning a half-water change was made and they were placed in the dark at 20°C. Bowls were then checked for egg bundles every half hour. Released egg bundles were distributed across 6-well culture plates for development in 15 PSU water in the 20°C incubator, in the dark. All gametes were incubated at the same temperature to minimize differences in larval thermal tolerance due to developmental plasticity or effects of temperature on developmental timing, allowing us to isolate parental effects as the cause of thermal tolerance differences. To clear gametes and promote gametogenesis during experimental conditions, parents were induced to spawn two weeks prior to the start of each experimental incubation.

### (f) Massachusetts parental effects experiment

To investigate if parental heat exposure could increase larval thermal tolerance, we compared larvae from STHS parents to those from control parents (figure 1B). Nine paired, genotype-controlled parental populations derived from the Massachusetts stock were used in this experiment. For six of the nine trials, parental populations were genetically identical across treatments, and for three of the nine trials, parental populations were genetically similar across treatments (i.e. there were unique lineages within treatments; see supplementary table 1).

Experimental populations were subjected to the STHS or control temperature regime and then cued to spawn as described above; larval thermal tolerance was assessed as described below. For each larval cohort 144-192 larvae were assessed. To test for the persistence of parental effects following STHS, three (genetically identical) parental population pairs were placed in the 20°C incubator after the initial STHS exposure and re-spawned after two weeks along with their paired control populations to account for any impacts in larval quality induced by repeated spawning.

### (g) Massachusetts maternal/paternal effects experiment

To investigate whether parental effects were predominantly paternal or maternal, six of the nine parental populations described above included additional female/male-only parental cohorts to enable reciprocal crosses of STHS and control males/females (figure 2A; table S1). Fertilization between STHS x control males/females was achieved by transferring eggs from female-only bowls into their opposite treatment male-only bowls. Female bowls were checked for the presence of new egg bundles every half hour, for five hours. For each larval cohort 144 larvae were assessed. In one lineage (G2), the STHS female x control male failed to produce viable larvae, and the STHS male x control female yielded only 96 viable larvae. For lineage G4, the STHS female x control male yielded only 48 viable larvae.

**Figure 2.**
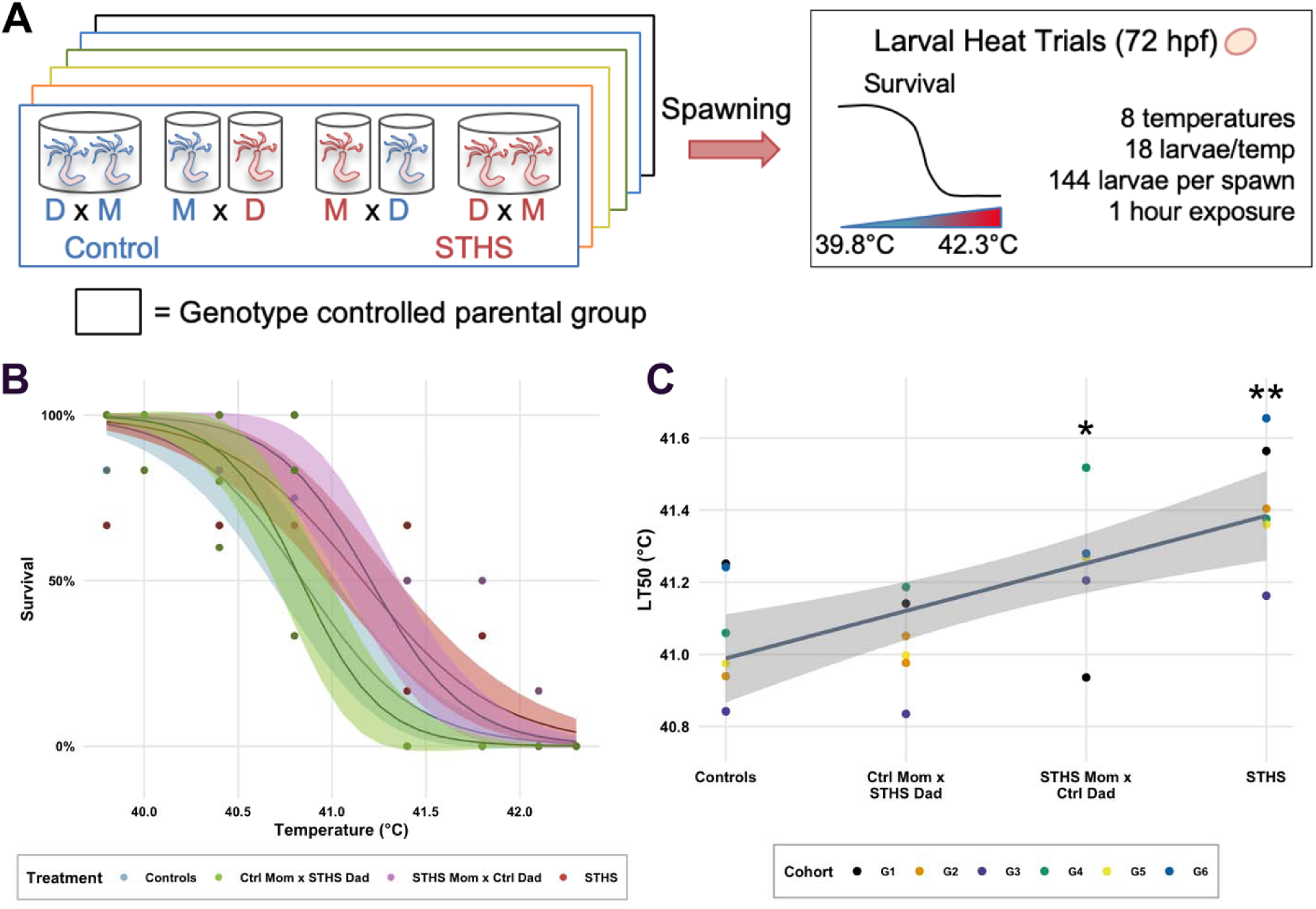
Increase in larval thermal tolerance is not attributable to exclusively maternal or paternal effects. (A) Experimental design. Colored boxes represent genotype-controlled parental populations. See table S1 for complete details. M: Mom; D: Dad (B) Survivorship curves of larvae from different parental treatments when exposed to acute temperature stress. Ribbon shows 95% confidence interval for logistic survivorship model. Curve shown for family G3, curves for all families tested are shown in figure S2. (C) LT50 for larvae from each family (N=6, colors), and parental treatment. Paired t-tests showed non-significant differences in LT50 of larvae from control parents and those with STHS father (*p*=0.37), and between controls and those with a STHS mother (*p*=0.14).

### (h) Local adaption vs. parental effects experiment (MA vs. NC)

To quantify how shifts in larval thermal tolerance from parental effects compared with adaptive differences between genetically and geographically distinct populations, we compared offspring from MA parents to those from NC parents. Due to the small number of unique NC genotypes, we used all genotypes to create a NC self-breeding population and to reciprocally cross with MA animals. Each parental combination had 14 females and 24 males. All four parental groups (MA, NC, MA female x NC male, NC female x MA male) were maintained under control conditions, spawned, and used to measure control larval thermal tolerance. After three weeks, the *same* parents were subjected to the STHS regime and re-spawned to measure larval thermal tolerance after parental heat exposure (figure 3A). Fertilization for MA x NC hybrids was conducted in the same manner described above for the maternal/paternal effects experiment. For each larval cohort 192-288 larvae were assessed for thermal tolerance.

**Figure 3.**
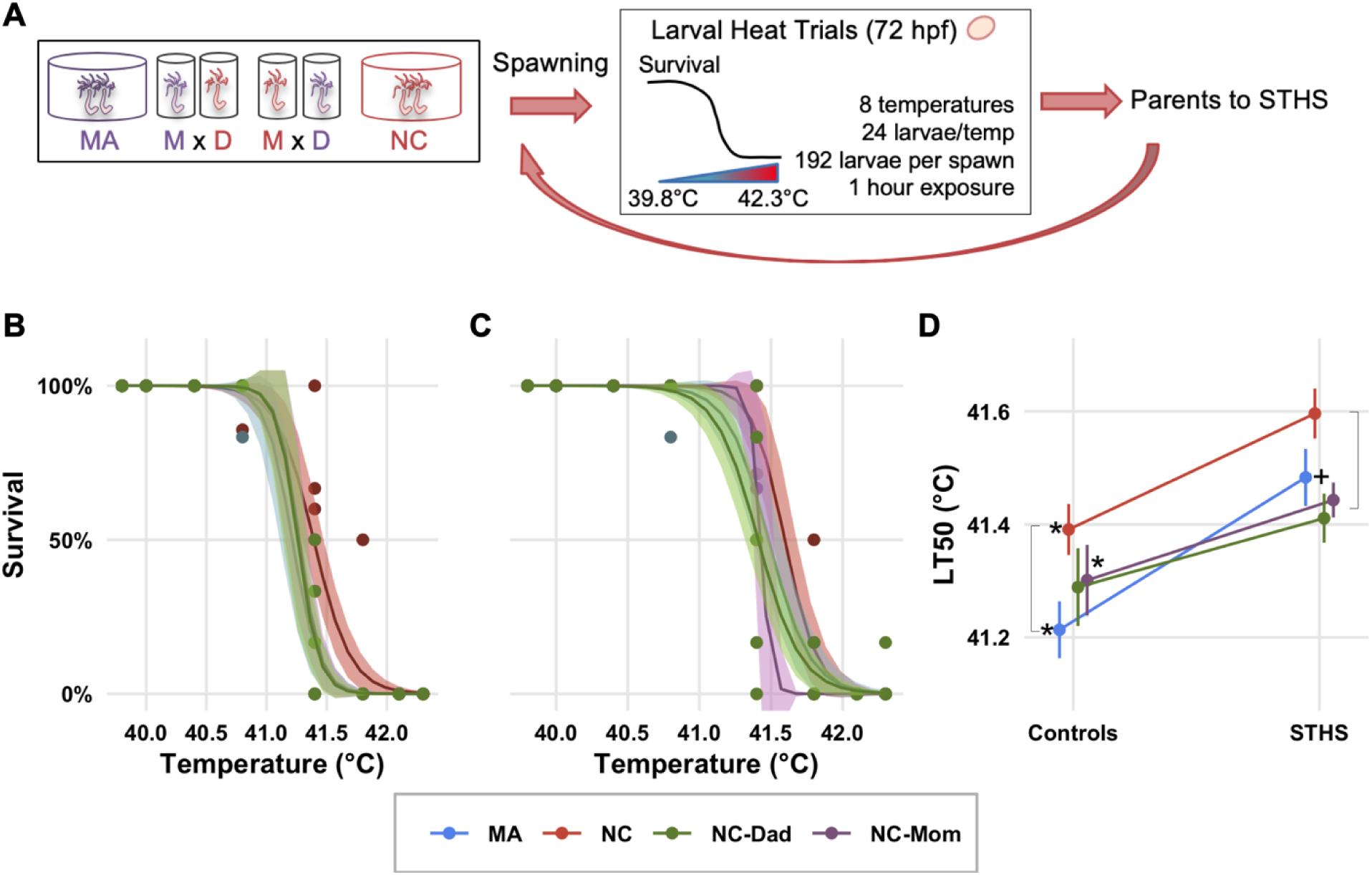
MA Parental effects confer thermal tolerance equivalent to native NC larvae. (A) Experimental design. Animals from NC are shown in red, from MA in blue. Note that unlike for MA-only experiments (figures 1-2), controlling for parental genotypes was achieved temporally instead of through the use of paired populations. The same groups of parents were spawned under control conditions and then after the STHS treatment. M: Mom; D: Dad (B) Survival of larvae under control parental conditions and (C) after parents underwent STHS ramp. (D) ΔLT50 for each parental cohort and cross. Asterisks denote populations for which there was a significant (*p*<0.05, likelihood ratio test) increase in larval LT50 following STHS exposure of parents. Brackets denote population pairs that had significantly different LT50 within parental treatment condition. Cross denotes *insignificant* difference between MA larvae from STHS parents and NC larvae from Control parents. LT50 of larvae from STHS MA parents is comparable to that of larvae from NC parents from both control conditions (*p*=0.17, likelihood ratio test) and STHS (*p*=0.09, likelihood ratio test).

### (i) Larval thermal tolerance assays

At 72 hours post fertilization (hpf), eggs that had developed into swimming planula larvae were individually pipetted into 0.2 mL PCR strip tubes (USA Scientific, Ocala, FL, USA) with 200 μl of 15 PSU water. Using a Bio-Rad™ C1000 PCR thermocycler (Bio-Rad Laboratories, Hercules, CA, USA), with two 48-well heating plates, larvae were exposed to a 39.8–42.3°C temperature gradient for one hour. Full protocol: (1) 1 minute at 25°C, (2) 4 minutes at 30°C, (3) 4 minutes at 38°C, (4) 1 hour at the treatment temperature: 39.8, 40, 40.4, 40.8, 41.4, 41.8, 42.1, or 42.3°C, (5) 4 minutes at 38°C, (6) 4 minutes at 30°C, and (7) infinite hold at 22°C. Each run consisted of six 8-well strip tubes (48 wells total) per larval cohort, such that larvae from treatment and control parents were always exposed simultaneously (one set per heating plate). The heating plate used for STHS and control larvae within each run was alternated to minimize any possible variations between the thermocycler’s heating plates.

Following the thermal exposure, larvae were maintained in the same PCR tubes and returned to the 20°C incubator, in the dark. Mortality was scored 48 hours after each trial by examining larvae under a dissecting scope. By this time, dead larvae had begun to disintegrate and appeared as fuzzy clumps. Surviving larvae were successfully maintained in PCR tubes for over a week, and showed metamorphosis during that time, suggesting mortality due to conditions in the tubes during the 48 hours between trials and scoring would be highly unlikely.

### (j) Statistical analysis of larval survival

A logistic regression model (LL.2) using the dose response curve, ‘drc,’ package (27) in R (28) was used to calculate survival curves and the dose (temperature) at which 50% of the larvae died (LT50) using the replicated sets for each larval cohort. In all experiments estimates for standard errors (of the LT50 estimate) were corrected for simultaneous inference using the glht() function from the ‘multcomp’ package in R. To compare the magnitude of change in LT50 values for larvae from STHS parents we analyzed experiments as follows:

#### MA parental effects experiment (including persistence)

The LT50 estimates from models for control and STHS parents were compared using paired t-tests to account for variation in thermal tolerance between family cohorts.

#### MA maternal/paternal effects experiment

The LT50 estimates were analyzed using a general linear model with parental treatment as a fixed effect and family as a random effect using the lmer() function from the ‘lme4’ package in R.

#### Local adaption vs. parental effects (MA vs. NC) experiment

For each pair of STHS and control curves (e.g. NC control, NC STHS) we tested whether response curves were significantly different by comparing a model in which both treatments were estimated to have the same LT50 and one in which LT50 estimate could vary by treatment, using likelihood ratio tests through the ‘anova()’ function in R. This method was used because there were a limited number of unique NC genotypes, therefore it would have been impossible to obtain independent replication at the level of parental groups between STHS and control treatments (see above).

### (k) Quantitative Polymerase Chain Reaction (qPCR) assays of larval gene expression

To test whether larvae from STHS parents exhibited differential expression of genes commonly involved in stress response pathways, we measured expression of three genes: Heat Shock Protein 70 (HSP70), Magnesium Superoxide Dismutase 2 (MnSOD2), and Citrate Synthase. HSP70 and MnSOD2 were chosen based on previous work on the response of *N. vectensis* to a variety of stressors, such as oxidative stress, UV, and pollutants (29,30). Citrate Synthase was chosen because activity is used as a proxy for mitochondrial density and capacity, which can play a role in thermal physiology (31,32). Four pairs of genetically identical parental populations were either subjected to the STHS or control thermal regimes (table S1). Larvae were allowed to develop as described above.

At 72 hpf, 2-5 replicate pools of 200-300 swimming planula larvae from each parental population and treatment were pipetted into 1.5 mL microcentrifuge tubes (figure 4A; table S2) for RNA extraction (“baseline” timepoint). Additional larvae were subjected to a 4-hour heat shock at 35°C using a Fisher Scientific™ Isotemp heating plate (Thermo Fisher Scientific, Waltham, MA, USA). Larvae were sampled for gene expression immediately after the heat shock (“immediate”) as well as 18 hours following the end of the heat shock (“post”) (table S2). For further details see supplementary methods.

**Figure 4.**
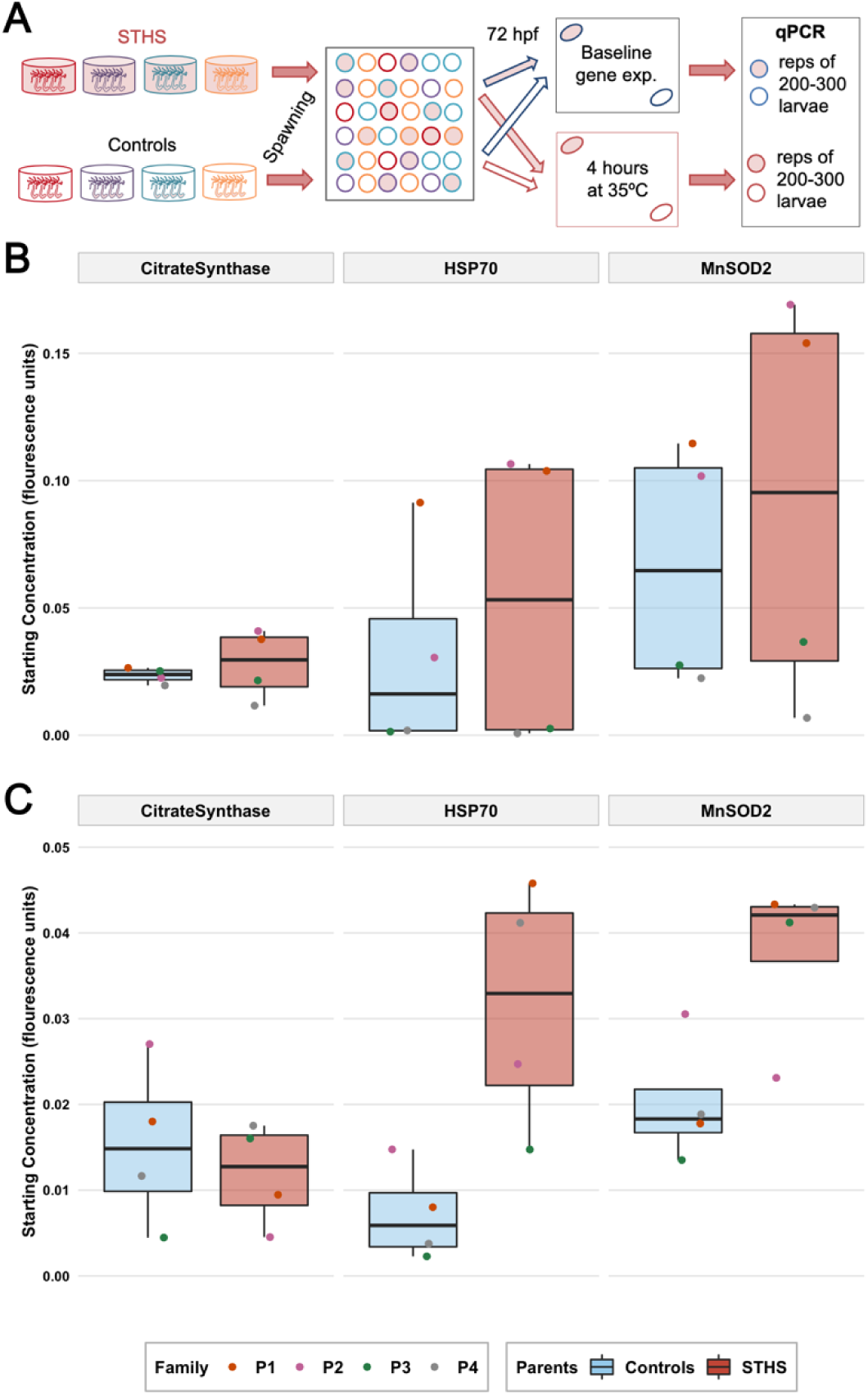
Larvae from STHS parents show elevated gene expression following heat stress. (A) Experimental design. Colored bowls represented genotype-controlled parental populations. Filled shapes indicate STHS parents and their larvae. (B) Constitutive gene expression at baseline in 72 hpf larvae from four control (blue) and STHS (red) clonal parental populations. Colored points correspond to parental population (family) (C) Inducible gene expression patterns 18 hours after the acute heat shock treat (bottom trajectory in panel A). Larvae from STHS parents a trend towards stronger induction of HSP70 and MnSOD2 following heat stress (*p=*0.052, *p=*0.12, respectively, paired t-tests). In both B and C expression is shown as the starting concentration after normalizing to expression of housekeeping genes and correcting for cross-plate variability (see supplementary methods).

For each gene, expression was measured in two 96-well plates with two technical replicates per sample, and two across-plate replicated samples, using Bio-Rad™ iTaq universal Syber Green Supermix, and a Bio-Rad™ CFX96 thermocycler (Bio-Rad Laboratories, Hercules, CA, USA) with the following protocol: (1) 1 min at 95°C, (2) 40 cycles of amplification [15 seconds at 95°C, 25 seconds at 60°C], and (3) a final melt curve from 65°C to 95°C with a 0.5°C increase every 5 seconds. Raw, uncorrected fluorescence values were used to estimate the starting template concentration using LinRegPCR (33). Cross plate variation was corrected using Factor_qPCR (34). To obtain final expression values, the geometric mean of the reference genes was used to normalize expression of each gene, by dividing the gene’s estimated concentration by the geometric mean of the references (see supplementary methods). Comparisons between values at different timepoints were conducted using paired t-tests.

### (l) Fertilized oocyte size measurements

To test whether the experimental conditions affected egg size and potential energetic provisioning, we examined egg bundles from all spawning females of four genetically identical parental population pairs from the MA parental effects experiment. We photographed eggs using a Zeiss™ Axio Cam 1Cc1 camera and imaging software (Carl Zeiss AG, Oberkochen, Germany). A stage micrometer was photographed under the equivalent magnification for scale. The diameter of 10 eggs per bundle was measured using Image J software (National Institutes of Health, Bethesda, MD, USA) (*N=* 230, 260, 160, and 220 eggs per control population and *N=*130, 240, 210, and 200 eggs per STHS population, respectively). Differences in mean diameters between STHS and control mothers for each population pair were tested using paired t-tests.

### (m) Adult heat stress survival assays

To test whether thermal preconditioning resulted in hardening of adult anemones to a subsequent heat shock, we exposed 80 mixed-genotype, MA adult individuals from the general laboratory populations to the STHS ramp described above and maintained another 80 individuals at 20°C. The day following the end of the STHS ramp, all anemones were subjected to a 6-hour heat shock at 36°C, and then returned to 20°C. Mortality was assessed 48 hours following the acute heat shock. Adults that had ejected mesenteries, were decomposing, or were unresponsive to touch were scored as dead. Difference in the proportion of surviving anemones by treatment was tested using a one-tailed, Fisher’s exact test in R for lower survival in control anemones.

## Results

### (a) Oocyte sizes before and after spawning

Immature oocytes were retained after spawning, with a significant shift in oocytes of smaller sizes following spawning, representing ∼45% decrease in size (*p*=3.07427×10^−5^, Wilcoxon test; figure 5B). Oocyte density was higher after spawning suggesting the release of larger, mature oocytes, and subsequent contraction of the mesenteries following spawning (mean: 21.8 oocytes/mm^2^ before spawning and 27.3 oocytes/mm^2^ after spawning). These data suggest selective spawning of mature oocytes, with retention of smaller immature oocyte that can continuously develop between spawns.

**Figure 5.**
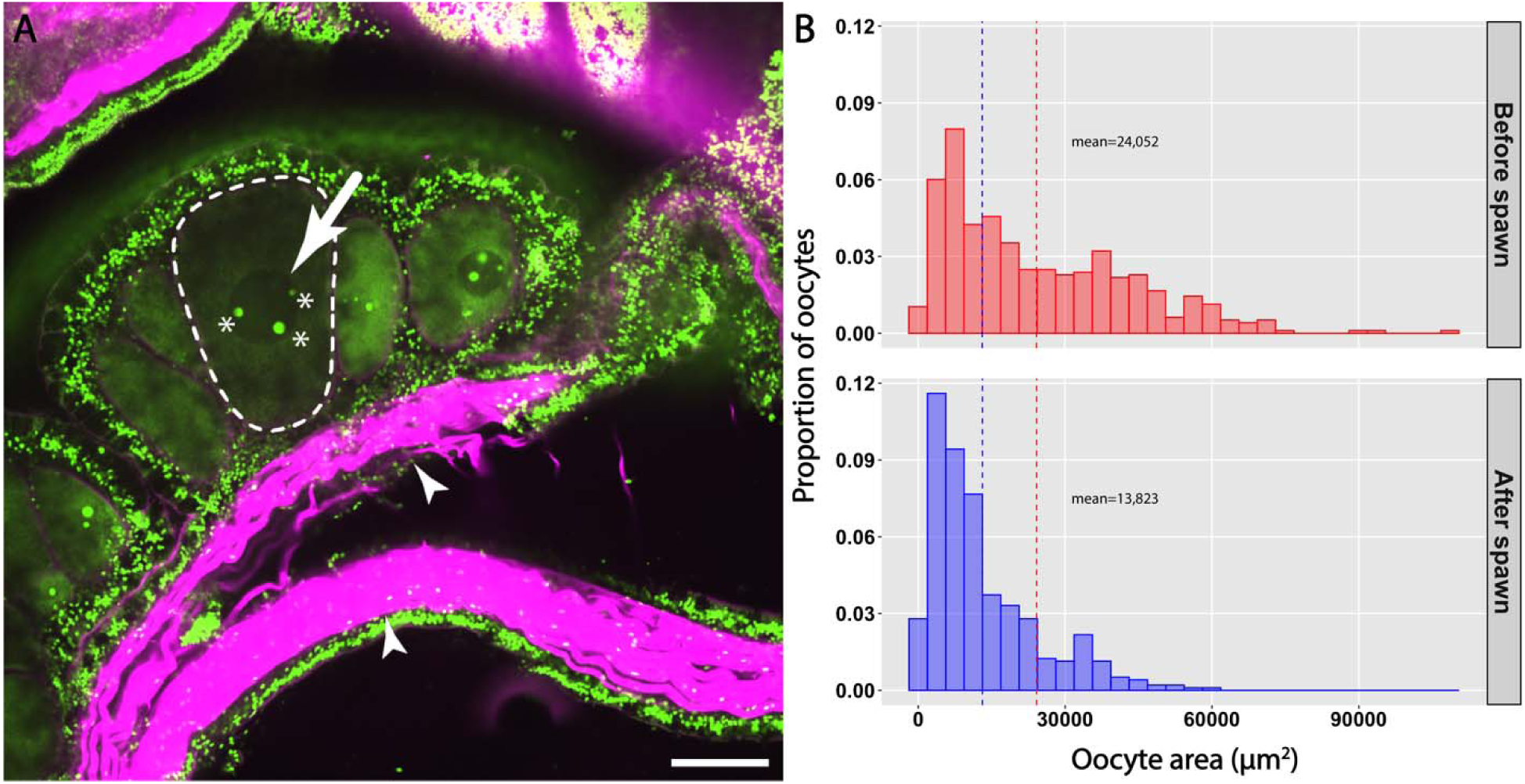
*Nematostella vectensis* females retain immature oocytes after spawning. (A) Single focal plane of confocal microscopy image of mesenteries containing oocytes. Oocytes (dashed outline) are distinguished from the surrounding tissue by size, the enlarged nuclei (white arrow) and nucleolus (asterisks). The retractor muscle fibers (white arrowheads) of the mesenteries are enriched with F-actin (*magenta*). Nuclei are labeled by SYBR™ Green I (*green*). Scale bar = 100 µm. (B) Distribution of oocyte area before and after spawn. After spawn, the oocytes remaining in gonads are significantly smaller (*p*=3.07427×10^−5^, Wilcoxon test). *Red* and *blue* dash lines denote mean oocyte size before and after spawn (24,052 and 13,823 μm^2^), respectively. *y*-axis is normalized oocyte count to total oocytes observed of each treatment (N=496 oocytes: before spawn; 469: after spawn).

### (b) MA parental effects

Across the nine clonal parental population pairs we found a significant (*p*=2.2×10^−7^, one-tailed, paired t-test) increase in LT50 (mean ΔLT50: 0.34°C, range: 0.22-0.46°C) in larvae from STHS parents (figure 1C-D). However, parental effects did not persist. Larvae from three parental cohorts that were re-spawned after 2 weeks no longer showed higher thermal tolerance (*p=*0.57, one-tailed, paired t-test; figure 1E).

### (c) MA Maternal vs. paternal effects

Linear model results indicate that maternal effects confer a significant (*p*<0.05) increase in larval thermal tolerance (mean ΔLT50: 0.18°C, SD: 0.087), while paternal effects induced no difference in larval LT50 compared to controls (mean ΔLT50: -0.001°C, SD: 0.084). Interestingly, larvae from parents that had *both* been subjected to STHS showed the largest increase in LT50 (mean ΔLT50: 0.369°C, SD: 0.082, *p*<0.01), indicating there are either synergistic effects to having both parents exposed to heat stress or epistatic effects when only the mother is exposed but not the father.

### (d) Genetic effects: North Carolina and Massachusetts crosses

As expected from previous studies, we found that NC purebred larvae had higher larval LT50 than MA purebred larvae under control conditions (ΔLT50: 0.18°C, *p=*0.008, likelihood ratio test, figure 3B-C). Hybrid larvae showed intermediate phenotypes, between MA purebred and NC purebred larvae (figure 3B-C). Exposure of MA parents to STHS, however, resulted in larvae that had statistically indistinguishable LT50 values to purebred larvae from NC controls (*p=*0.17, likelihood ratio test), and from NC STHS larvae (*p*=0.09, likelihood ratio test).

NC purebred larvae from STHS parents also showed a significant increase in larval LT50 compared to the controls (ΔLT50: 0.2°C, *p*=0.001). Hybrid larvae with a NC mother and a MA father showed a significant, though smaller increase following STHS (ΔLT50: 0.17°C, p=0.001). Hybrids from a MA mother and NC father showed a trend towards higher LT50, but this shift was not statistically significant from controls (ΔLT50: 0.13°C, *p*=0.09).

### (e) Larval gene expression

Spawning success across parental cohorts used for this experiment was uneven, therefore, replication depth across different parental groups and larval treatments varied (electronic supplementary material, table S2). Overall, gene expression levels did not differ between larvae from STHS and control parents at baseline conditions (*p=*0.29 for citrate synthase, *p=*0.31 for HSP70, and *p=*0.25 for MnSOD2; paired t-tests). Eighteen hours following the acute larvae heat stress, larvae from STHS parents showed a trend toward higher expression of HSP70 (*p=*0.052, paired t-test), and MnSOD2 (*p=*0.12, paired t-test), but not Citrate Synthase (*p=*0.68, paired t-test) (figure 4C).

Gene expression was also measured immediately following the larval heat stress for larvae from control parents across all families and in larvae from STHS family P2. Expression of HSP70 was significantly higher immediately following heat stress in larvae from control parents relative to baseline (*p=*0.03) while expression of Citrate Synthase was significantly lower (*p=*0.04, respectively, paired t-test; figure S4). Gene expression results across all families and time points are shown in figure S5.

### (f) Fertilized oocyte sizes

We did not find any differences in fertilized oocyte diameter by control or STHS mothers (*p*=0.28, paired t-test) (figure S3).

### (g) Pre-conditioning of adult anemones to heat stress

We found a significant difference (*p*=0.03, Fisher’s exact test) in survival following an acute, 6-hour heat shock at 36°C between STHS and control adult anemones. On average there was 80% survival in control anemones and 92% survival in STHS anemones, with an odds ratio of 0.38, indicating anemones exposed to the STHS regime are less likely to die from an acute heat shock. This is what would be expected if thermal exposure induced physiological hardening in *N. vectensis.*

## DISCUSSION

### (a) Parental effects substantially increase larval thermal tolerance and respond rapidly to changing environments

In the context of global climate change, phenotypic plasticity has emerged as an avenue for organisms to persist through rapid environmental change, while the “slower” mechanisms of selection and genetic (mutation-based) adaptation catch up (35–38). The degree to which such plasticity can benefit organisms will depend on species- and environment-specific interactions (see (39–41) for such considerations). Despite caveats, parental effects and transgenerational plasticity – usually epigenetic or other semi-heritable changes across generations – are being heralded as a potential safety net for vulnerable species.

We used *N. vectensis* as a powerful study system to examine the relative strength of phenotypic plasticity in parental effects compared to adaption in modulating larval thermal tolerance. We found that thermal threshold of larvae from heat-exposed Massachusetts parents was comparable to that of larvae from control North Carolina parents (figure 3B-C) – a genetically distinct population, locally adapted to warmer temperatures (18). This demonstrates parental effects can confer rapid shifts in larval thermal tolerance that are ecologically relevant and of a similar magnitude to shifts driven by adaption. Heat exposure of North Carolina parents also resulted in increased larval thermal tolerance (figure 3B-C), signifying responsive parental effects are not unique to Massachusetts populations, and may, instead, be an inherent characteristic of *N. vectensis* that suits its naturally variable estuarine habitat. The shallower slope in the reaction norm of hybrid larvae, however, suggests that negative epistatic effects may arise when combining genotypes between populations and that differences in larval thermal tolerance between sites have a strong genetic basis.

Controlling for genetic variability, through the use clonal lineages, offered greater power to detect shifts in thermal tolerance, as we found substantial differences between larvae from different parental cohorts under both control and STHS parental conditions (figures 1C-E, 2B-C). For instance, LT50 estimates for larvae from control parents ranged from 40.84 to 41.25°C, while those for larvae from STHS parents ranged from 41.16 to 41.65°C. Remarkably, however, the induced change in larval thermal tolerance following exposure of parents to STHS was highly consistent across parental lineages (mean=0.34°C ±0.07 SD), suggesting *N. vectensis* show a consistent parental response to environmental variation. Whenever possible, studies should control for parental genotypes in order to increase sensitivity in detecting subtle changes in organismal responses.

Despite the increasing popularity of studies in parental effects and transgenerational plasticity, many studies only test the phenotypes of one cohort of offspring. For organisms with multiple reproductive cycles it is important to also test the persistence of such effects across subsequent breeding periods. In our case, we find that *N. vectensis* quickly readjusts after the heat regime ends. Larvae from the parents’ next spawn return to baseline levels of thermal tolerance (figure 1E), indicating reversible (plastic) parental effects in *N. vectensis*. The continuous gametogenesis cycle in *N. vectensis* likely facilitates this process by maintaining a constant pool of immature oocytes that can be modified by cues from the environment in which they directly develop (figure 5). Given the anemone’s naturally fluctuating environment, this plasticity in parental effects combined with continual gametogenesis could be more beneficial than adjusting irreversibly in any one direction (42). Our results underscore the need to test for reversibility or persistence across breeding periods, as well as monitoring of gametogenesis, in order to better understand how such mechanisms may help (or fail to help) organisms keep pace with global climate change.

It is also important to determine what degree of parental exposure or stress yields beneficial results for offspring. In Massachusetts, *N. vectensis* experience a wide diel (approximately 15°C during summer months), and seasonal range in temperatures (approximately 40°C between winter and summer temperatures). In pilot studies (data not shown), we tested a longer parental exposure condition that kept parents at 30°C for four weeks. Offspring from those parents still showed higher thermal tolerance than controls, but the effect was smaller than the observed increase in thermal tolerance following the STHS regime, suggesting chronic stress exposure may limit the ability of parents to adequately provision larvae. In addition, experiments that use variable temperature conditions that more accurately reflect *N. vectensis*’s natural environment may show different results than those using static temperature conditions.

### (b) Larval thermal tolerance is influenced by maternal and combined parental effects

We find exposure of mothers alone to the STHS regime leads to an increase in larval LT50, while exposure of fathers alone does not result in any detectable shift in larval thermal tolerance (figure 3C). Maternal effects alone, however, account for only about half of the LT50 increase seen when both parents are exposed to the STHS regime, suggesting that having both parents under the same conditions confers additional benefits (figure 3C). The mechanisms responsible this synergy are unknown but theoretical epistatic effects between maternal effects and offspring genotypes have been described (43).

While examples of maternal effects abound, direct paternal effects have been described in only a few species (44). Nevertheless, paternal influences on zygotic phenotypes through transfer of cytosolic compounds to the fertilized egg or through compounds in the seminal fluids have been described in humans (45), trout (46), mice (47), and insects (48). It is possible that male *N. vectensis* are providing larvae with additional resources but these are insufficient when decoupled from maternal contributions. Potential influences of maternal effects on the efficacy of paternal effects have also been conceptually explored (44), and have been shown for paternally inherited QTLs that influence thermal tolerance in rainbow trout (46). These patterns, combined with the elevated expression of HSP70 and MnSOD2 in larvae from STHS parents following heat stress, hint that provisioning effects such as RNA loading may contribute to larval thermal performance.

### (c) Potential mechanisms of increased larval thermal tolerance through parental effects

Uncovering the mechanisms behind these effects can also improve understanding of organismal physiology under changing conditions. Preliminary experiments (data not shown) showed a substantially smaller increase in thermal tolerance for juveniles (7 dpf) from STHS parents, suggesting that gamete provisioning mechanisms (such as increased lipid or antioxidant content, or transcript loading) may be more likely than effects that last further into development (e.g. induction of developmental plasticity), as these would enhance fitness of early larval stages but wane as larvae grow. The timing of gametogenesis in relation to stress exposure may also play an important role, as has been shown for polychaetes (8). During our trials, we spawned animals prior to the start of the experimental heat ramps, which facilitates clearance of mature oocytes (figure 5B). This would imply that the experimental larvae were primarily derived from oocytes that completed later stages of maturation in the experimental conditions. However, we cannot be certain that all the mature gametes were released during the pre-spawn. It is possible, therefore, that some of the effects described here arise from a combination of direct parental effects and environmental effects on developing gametes, as the timing of gametogenesis cannot be entirely constrained to the experimental period (see (36) for a detailed discussion of such considerations). We investigated two possible mechanisms for increased larval tolerance: modulation of egg size and of larval gene expression. We found eggs from STHS and control parents did not differ in size. However, because we did not characterize the biochemical composition of the eggs, it remains possible that the availability of specific resources, such as lipids or antioxidants differed between parental treatments. In regard to gene expression, larvae from STHS parents showed stronger induction of HSP70 and MnSOD2 expression eighteen hours after heat stress, suggesting that sustained expression could be influencing tolerance (figure 4C). However, larvae from control parents, showed a prompt return to baseline (pre-stress) levels of gene expression after showing differences in expression immediately after heat stress (figure S4). This pattern, termed ‘transcriptomic resilience,’ has been linked to stress tolerance in other species such as seagrasses and corals (49,50). As such, it is surprising that transcriptomic resilience in larval gene expression patterns for *N. vectensis* is not coupled with higher thermal tolerance (i.e. larvae from control parents tend this pattern, not larvae from STHS parents). In particular, whether a gene was up- or down-regulated differed substantially by family, suggesting differences in gene expression dynamics by familial cohort. If gene expression is contributing to larval tolerance, then it is possible that the dynamics of expression (e.g. timing and magnitude) are more important than either factor alone. Due to limitations in sampling size, our study may have missed other dynamic expression patterns or lacked the power to detect substantial differences in expression across timepoints. Future transcriptomics studies could reveal more complex patterns and identify other candidate genes involved in increasing larval thermal tolerance.

Exposure of *N. vectensis* to elevated temperatures also substantially increased survival of adult anemones to acute heat shock, suggesting *N. vectensis* can also modulate adult physiology to match thermal conditions. A recent study of adult thermal acclimation to a range of temperatures (6 - 33°C) suggests *N. vectensis* rapidly adjusts respiration rates and later increases metabolic capacity (activity of mitochondrial enzymes) when exposed to different temperatures (Brinkley, Rivera, and Tarrant, unpublished data). Studies that focus on mitochondrial physiology in adults and larvae may help identify the mechanisms responsible for *N. vectensis’* rapid acclimation to different thermal regimes.

While plasticity both of adult physiology and parental effects on their offspring can enhance short-term survival, there may also be trade-offs. For instance, heritable transgenerational changes that increase offspring’s aerobic scope have been described in spiny damselfish (51), but F2 generation fish maintained at warmer temperatures were unable to breed, suggesting a strong trade-off between thermal performance and reproduction (52). Such trade-offs should be explored in future studies in order to fully characterize the potential of parental effects to promote species persistence under changing climate scenarios.

### (d) Conclusions

We find that parental effects on thermal tolerance in *N. vectensis* are short-lived and quickly reversible (subsequent cohorts lose protection) suggesting *N. vectensis* responds quickly to its current environment and may take advantage of parental effects without long-term trade-offs. Studies that follow multiple generations of *N. vectensis* through parental heat exposure and track offspring growth and eventual reproduction could elucidate any potential trade-offs associated with increased thermal tolerance early in life. A parental strategy that favors short-term gamete provisioning over longer-term epigenetic changes may be better suited to *N. vectensis’* highly variable yet broadly predictable (seasonal and tidal) environment.

Studies of *N. vectensis* reproduction in the field are scarce. Only one study, to our knowledge, describes gravid gonads in field-collected individuals from Nova Scotia, and only during August and September (53). A handful of other studies suggest reproduction is mainly asexual under natural conditions, given the high levels of clonality observed in field collected anemones (16,54,55). Dedicated studies of *N. vectensis’* reproductive cycle in the field would offer greater understanding of its parental strategies.

Overall, *N. vectensis* offers a robust organismal system in which to study thermal responses due to their wide thermal range, easy husbandry, fast development, ease of spawning, ability to generate clonal lineages, as well as their well-developed genomic resources. Here we show *N. vectensis* is capable of quickly modulating parental effects to increase larval thermal tolerance. While maternal exposure can result in significant shifts, exposure of both parents to different environments results in a stronger shift in larval thermal tolerance. In a northern population (MA), shifts due to parental effects result in larval thermal limits that are comparable to those of a southern, more thermally tolerant, population (NC). These patterns point to both a genetic and plastic basis for thermal tolerance in *N. vectensis*. Given our rapidly changing global thermal environment, studies that aim to uncover the mechanisms responsible for these rapid shifts in thermal performance can provide insights into the sensitivity, acclimation, and adaptation potential of vulnerable species such as marine ectotherms.

## Data accessibility

Additional methods and results as noted in the paper, are in the electronic supplementary material. Raw data and code for all statistical analyses and figure generation are included in the authors’ GitHub repository: https://github.com/hrivera28/Nematostella-ParentalEffects/ and https://github.com/Penguinayee/Nematostella_ParentalEffects

## Author contributions

H.E.R and A.M.T. conceived of the experimental design, maintained laboratory populations, and conducted experiments. H.E.R. generated clonal lineages, analyzed the data, and wrote the initial draft. C.Y.C. and M.C.G. maintained laboratory populations used for oocyte analyses. C.Y.C. sampled, prepped, imaged anemones, and analyzed oocyte data. All authors read and approved the final manuscript.

## Acknowledgements

We would like to thank Rebecca Helm for initial guidance on experimental questions and design, Victoria Starczak for guidance on statistical analyses, and Hollie Putnam for input on appropriate experimental designs to investigate the role of parental and transgenerational effects. We further thank Hadley Clark, Sabine Angier, and Hannah Stillman for their assistance with animal care and larval trials. We also thank Adam Reitzel and for providing animals from North Carolina and Whitney Leach for field assistance. We thank David Brinkley, Jehmia Williams, Sarah Davies, Hannah Aichelman, Nicola Kriefall, Daniel Wuitchik, and James Fifer for feedback on figures. We also thank members of H.E.R.’s thesis committee (Iliana Baums, Simon Thorrold, Janelle Thompson, and Anne Cohen) for feedback on data analysis and interpretation. Lastly, we thank the Cnidofest 2018 meeting for facilitating collaborations between the Tarrant and Gibson labs.

## SUPPLEMENTARY METHODS

### Larval qPCR methods

At 72 hpf, 2-6 replicates of 200-300 swimming planula larvae from each parental population and treatment were pipetted into 1.5 mL microcentrifuge tubes (table S2). Tubes were spun down for 10 seconds to concentrate larvae and 15 PSU water was removed until only 100 μl of water remained.

All larvae were immediately processed for RNA extraction using the phenol-chloroform based Bio-Rad™ Aurum Total RNA Fat and Fibrous Tissue Kit (Bio-Rad Laboratories, Hercules, CA, USA) with on-column DNase treatment. RNA yields were assessed using a NanoDrop One™ spectrophotometer, Thermo-Fisher™ (Thermo Fisher Scientific, Waltham, MA, USA), giving a mean yield of 44.9 ng/μl and a range of 12.8-132.2 ng/μl in a 15 μl total elution volume. For synthesis of complimentary DNA (cDNA) we used 200 ng of RNA per sample and the Bio-Rad™ iScript DNA Synthesis kit and Bio-Rad™ C1000 PCR thermocycler (Bio-Rad Laboratories, Hercules, CA, USA) with the following protocol: (1) 5 minutes at 25°C (2) 20 minutes at 46°C (3) 1 hour at 95°C, and (4) 4°C hold.

Primer sequences for Actin, 18S, L10, HSP70, and MnSOD2 were obtained from (26) and (27) as these were previously used in *N. vectensis*. The gene sequence for Citrate Synthase was determined by searching the *N. vectensis* genome on the JGI and selecting the eukaryotic-type CS annotated sequence. The full sequence was then submitted to the Primer3 web portal to generate the best primer sequence. Synthesized primers were obtained from Thermo Fisher Scientific (Waltham, MA, USA). Actin, 18S, and L10 were used as reference genes. Gene accession numbers and primer sequences are listed in table S3.

## SUPPLEMENTARY FIGURES

**Figure S1.**
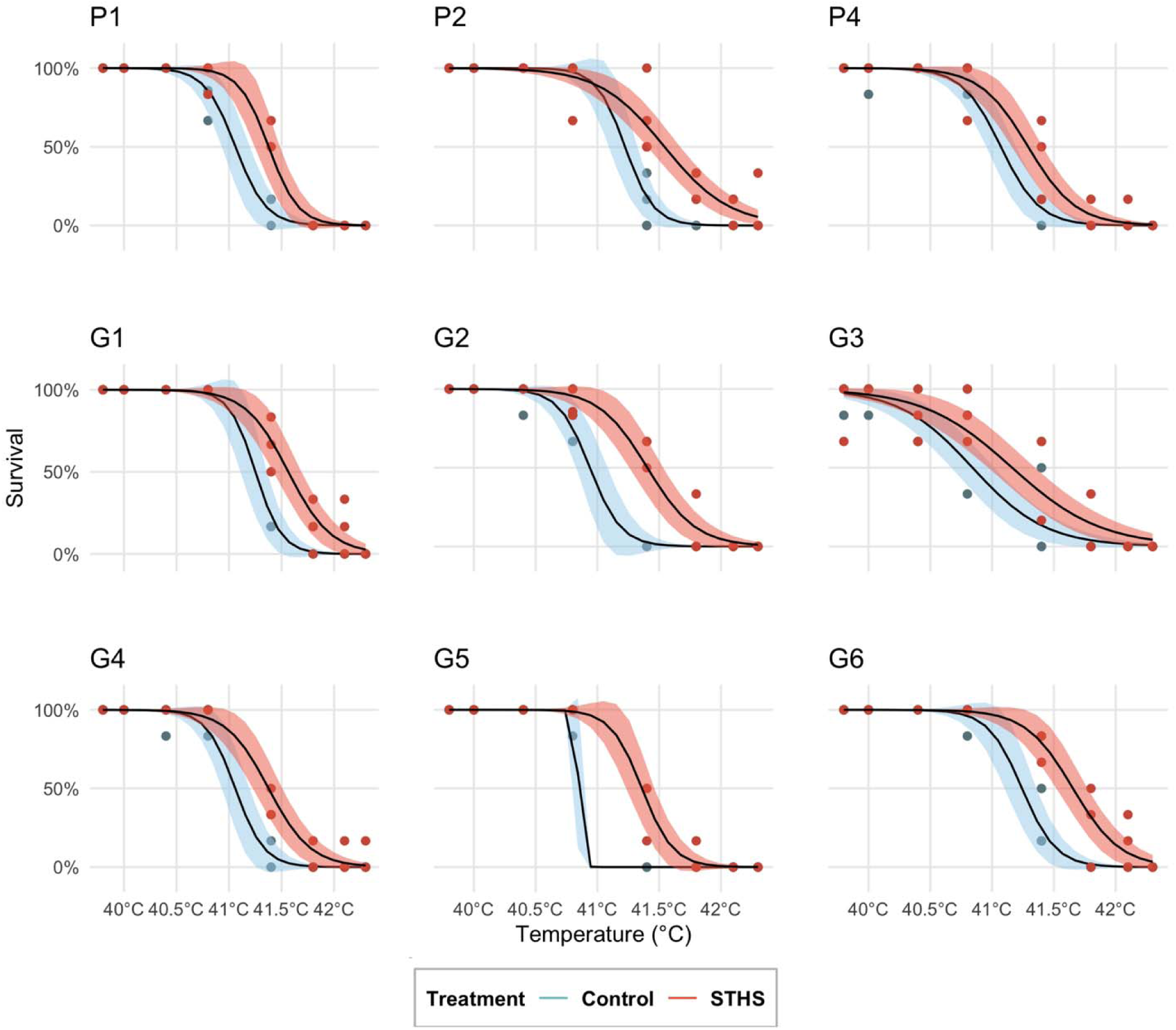
Survival curves for all larval cohorts from MA parental effects experiment. Survival of larvae from different parental treatments when exposed to acute temperature stress. Ribbon shows 95% confidence interval for logistic survivorship model. Parental population codes correspond to populations described in table S1.

**Figure S2.**
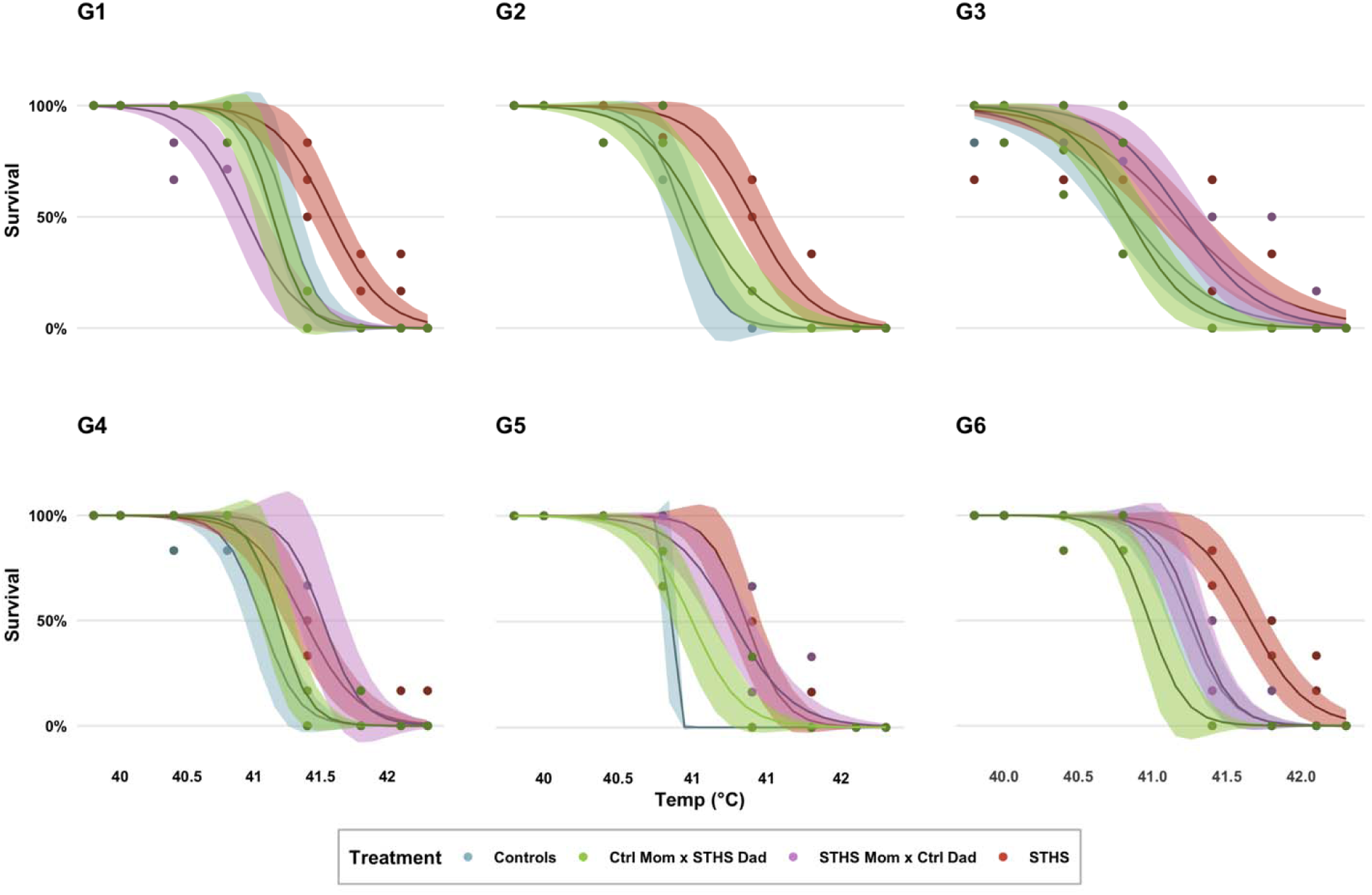
Survival curves for all larval cohorts in MA maternal/paternal effects experiment. Survival of larvae from different parental treatments when exposed to acute temperature stress. Ribbon shows 95% confidence interval for logistic survivorship model. Parental population codes correspond to populations described in table S1.

**Figure S3.**
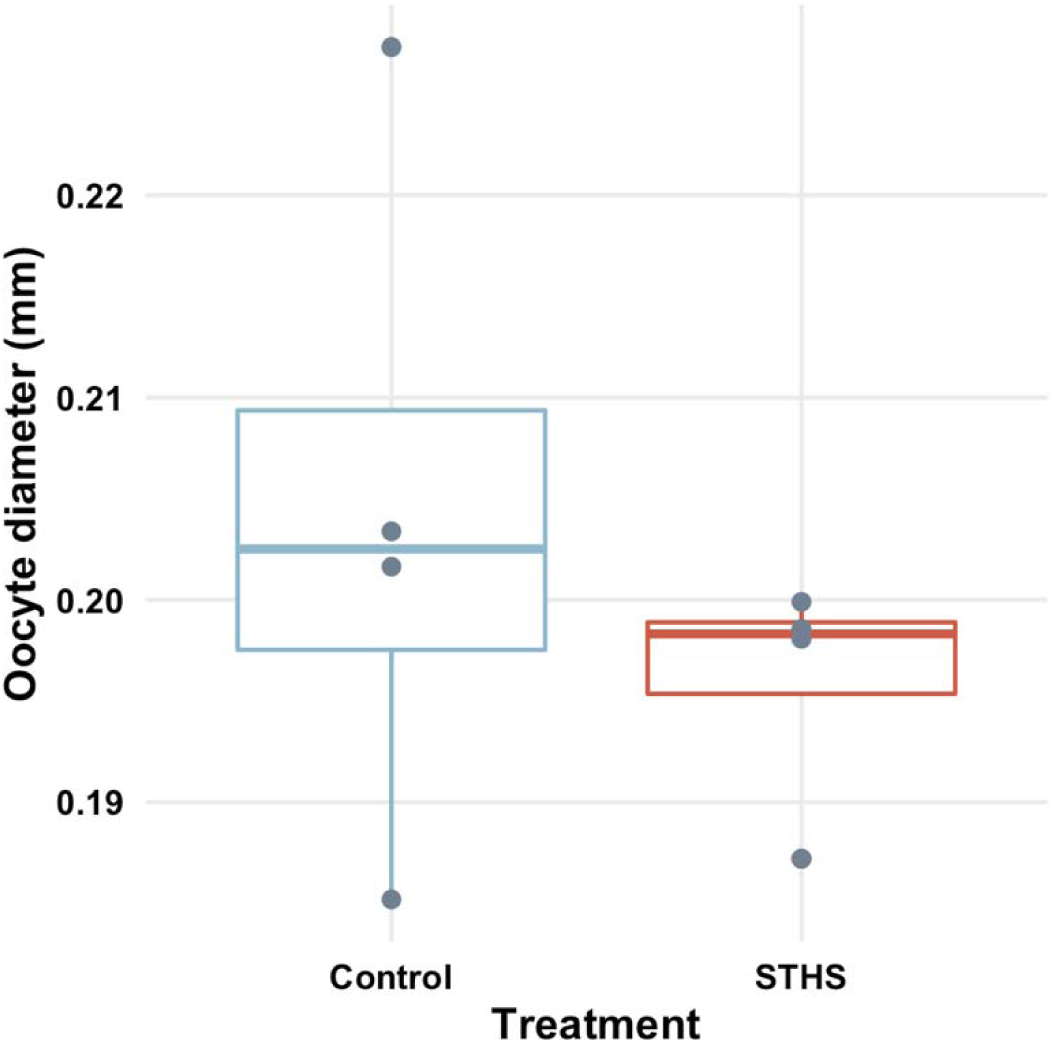
Egg diameter does not differ between parental treatments. Diameter of eggs released by four genetically identical paired parental populations. There is no significant difference in sizes of eggs by parental treatment (*p*=0.28, paired t-test).

**Figure S4.**
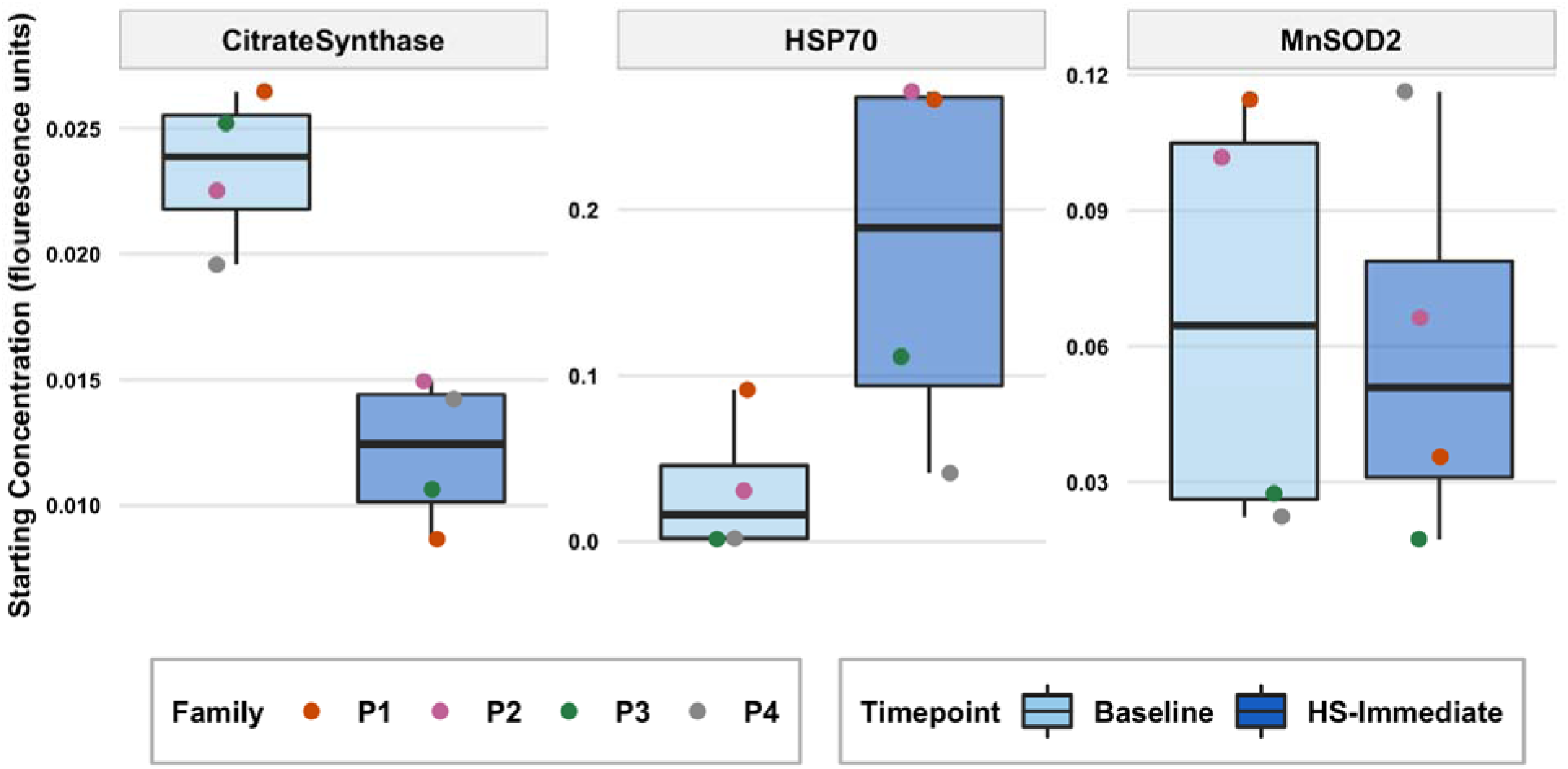
Larvae from control parents show differences in expression immediately after heat stress. Gene expression is shown as the starting concentration after normalizing to expression of housekeeping genes and correcting for cross-plate variability (see methods). Baseline: Constitutive expression at 72 hpf without any heat exposure. HS-Immediate: Immediately following the acute larval heat shock. There was significantly lower expression of citrate synthase after heat stress (*p*=0.03, paired t-test), but significantly higher expression of HSP70 (*p*=0.04, paired t-test).

**Figure S5.**
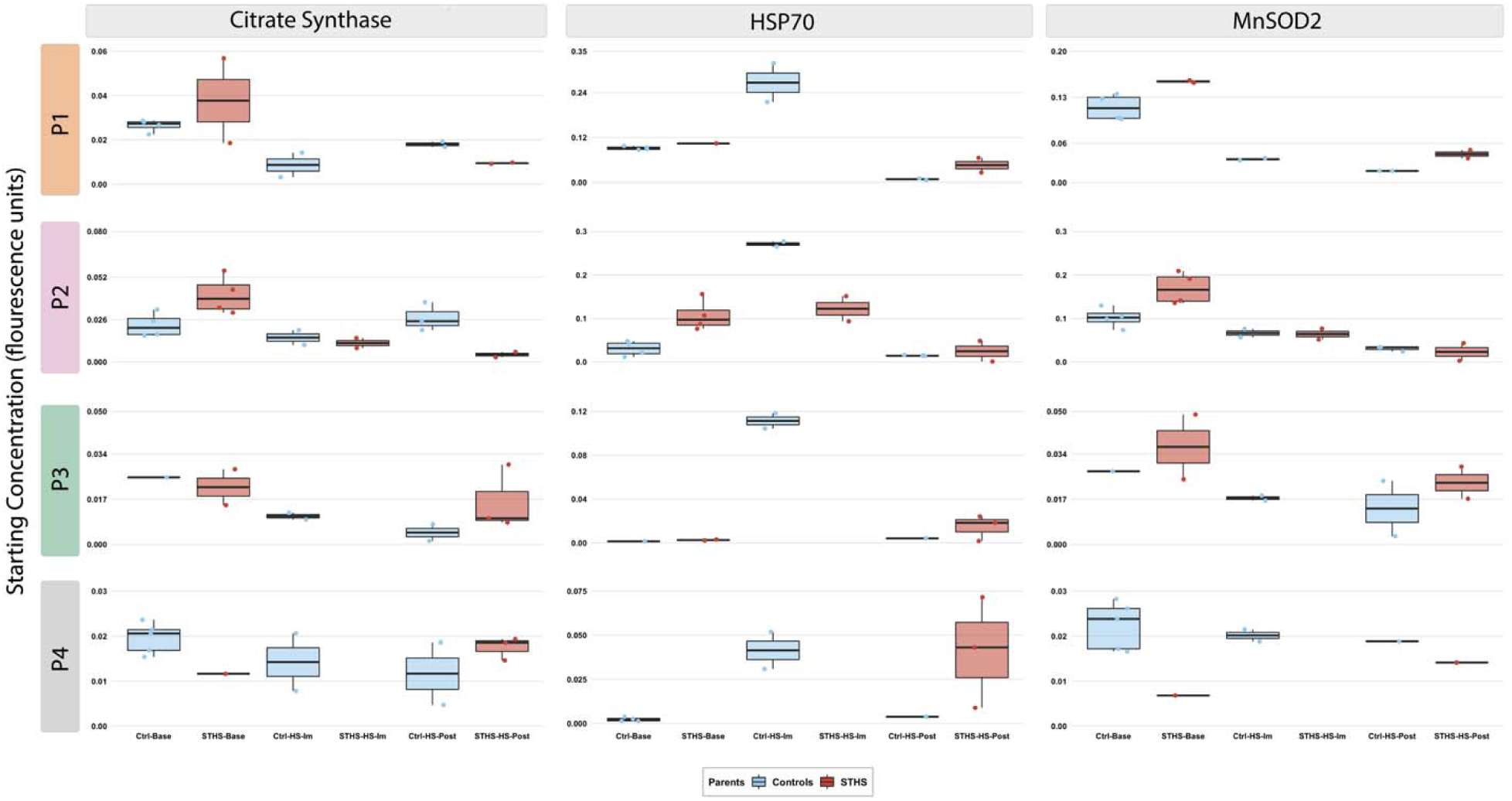
Larval gene expression across all timepoints and families. Gene expression is shown as the starting concentration after normalizing to expression of housekeeping genes and correcting for cross-plate variability (see methods). Base: Constitutive expression at 72 hpf without any heat exposure. HS-Im: Immediately following the acute larval heat shock. HS-Post: 18 hours following the end of the heat shock. Larvae from control parents are designated with “Ctrl” and those from STHS parents with “STHS.” Expression is faceted by family (rows) and genes (columns).

